# Hybrid yak-cattle *in situ* conservation via interspecies somatic cell nuclear transfer at ultra-high-altitude region

**DOI:** 10.1101/2025.07.10.664179

**Authors:** Dawei Yu, Qunzong Nima, Lei Cao, Shigang Gu, Kangjun Yin, Yurong Zhang, Jing Wang, Wangdui Basang, Yongye Huang

**Affiliations:** State Key Laboratory of Animal Biotech Breeding, Institute of Animal Science, Chinese Academy of Agricultural Sciences, Beijing 100193, China; National Germplasm Center of Domestic Animal Resources, Institute of Animal Science, Chinese Academy of Agricultural Sciences, Beijing 100193, China; Xizang Autonomous region Animal Husbandry General Station, Lasa Xizang 850000, China; Institute of Animal Husbandry and Veterinary Medicine, Tibet Academy of Agriculture and Animal Husbandry Sciences, Lhasa 850009, China; State Key Laboratory for Germplasm Resources and Genetic Improvement of Highland Barley and Yak Jointly Constructed by the Ministry and the Province, Lhasa 850009, China; College of Life and Health Sciences, Northeastern University, Shenyang 110169, China

**Keywords:** somatic cell nuclear transfer, cattle, yak, high-altitude, embryo, *in situ* conservation

## Abstract

Pien-niu, the hybrid offspring of yak (Bos grunniens) and domestic cattle (Bos taurus), possess exceptional adaptability to ultra-high-altitude environments of the Qinghai-Xizang Plateau. *In situ* conservation is of special significance for ultra-high-altitude region because the offsprings need to possess the ability to adapt to the unique environment. The present study was to clone Pien-niu in Xizang via interspecies somatic cell nuclear transfer (iSCNT) to practice *in situ* conservation of large animal at ultra-high-altitude region. Ear fibroblast cells were isolated from Pien-niu at Qushui Experimental Station (3650m above sea level), an animal live conservation farm at Xizang, and used as iSCNT donor cells. The iSCNT blastocysts were transferred into the oviduct of surrogate Pien-niu which are maintained at Qushui Experimental Station. A live cloned Pien-niu calf was born on May 12, 2025 and keeps healthy till now. Results of short tandem repeat analysis revealed that the microsatellite loci of cloned calf were completely matched with that of donor fibroblast cells. The successful cloning of Pien-niu can provide a paradigm for *in situ* germplasm conservation via iSCNT at ultra-high-altitude region.

## 1. Introduction

The hybrid offspring (Pien-niu) produced by crossing yaks (Bos grunniens) with domestic cattle (Bos taurus) not only retain the yak’s exceptional adaptability to ultra-high-altitude environments but also exhibit superior meat and milk production traits inherited from cattle. However, the prevalent male sterility of Pien-niu ^1^ severely restricts the utilization of this hybrid advantage in genetic resource development. Somatic cell nuclear transfer (SCNT) technology offers a potential solution for the asexual propagation of superior Pien-niu individuals. Since the first cloned cattle were produced in 1999, this technique has successfully generated multiple physiologically normal cloned cattle in low-altitude regions ^2-4^. This study aims to break through technical barriers and achieve the first animal cloning in the ultra-high-altitude environment of Xizang Autonomous Region, China. Notably, existing research indicates that prolonged exposure to hypoxia may lead to impaired reproductive function and abnormal placental development in animals ^5,6^. It suggests that the unique extreme environmental factors of the Qinghai-Xizang Plateau, including hypobaric hypoxia and intense radiation, may pose significantly greater challenges for cloning technology implementation compared to low-altitude areas.

Interspecies somatic cell nuclear transfer (iSCNT) demonstrates distinct advantages over conventional SCNT techniques ^7,8^, offering novel approaches for germplasm resources conservation and therapeutic cloning, albeit with significantly increased technical complexity. To date, this technology has been successfully applied in multiple mammalian species, including the African wildcat (Felis silvestris lybica)^9^ and European mouflon (Ovis orientalis musimon) ^10^. To expand the potential applications of cloning technology in biotechnological breeding and germplasm resources conservation, collecting cattle ovaries from the slaughter nearby to generate iSCNT recipients for cloning Pien-niu would be more feasible.

## 2. Results

### 2.1 Generation of iSCNT embryos using Pien-niu ear fibroblast cells as donor

To conduct iSCNT, ear fibroblast cells were harvested from a Pien-niu (Figure 1) at Qushui Experimental Station and used as donor cells. The iSCNT recipients were *in vitro* maturated cattle oocytes which were derived from ovaries collected from a slaughter nearby. As shown in Table 1, the cleavage and blastocyst formation rates reached 67.7±13.4% and 34.9±2.9%, respectively. The cleavage rate was lower than that in SCNT embryos using cattle ear fibroblast cell as donor. The blastocyst formation proportion was lower than both SCNT and *in vitro* fertilization (IVF) embryos. However, it should be noted that the iSCNT reconstructed blastocysts exhibited normal morphology. Well-developed blastocysts were selected to be used for embryo transfer.

**Table 1.** Assessment of *in vitro* development of SCNT embryos.

| Type | Donors | Recipients | No. of<br>reconstructed<br>embryos | % cleaved<br>± S.E.M. | % blastocyst<br>± S.E.M. |
| --- | --- | --- | --- | --- | --- |
| SCNT | Pien-niu ear<br>fibroblast cell | Cattle oocytes | 227 | 67.7±13.4 | 34.9±2.9 |
|  | Cattle ear fibroblast<br>cell | Cattle oocytes | 199 | 91.8±1.8 | 40.8±10.9 |
| IVF | Yak sperms | Cattle oocytes | 201 | 71.8±11.2 | 41.5±6.6 |

**Figure 1.**
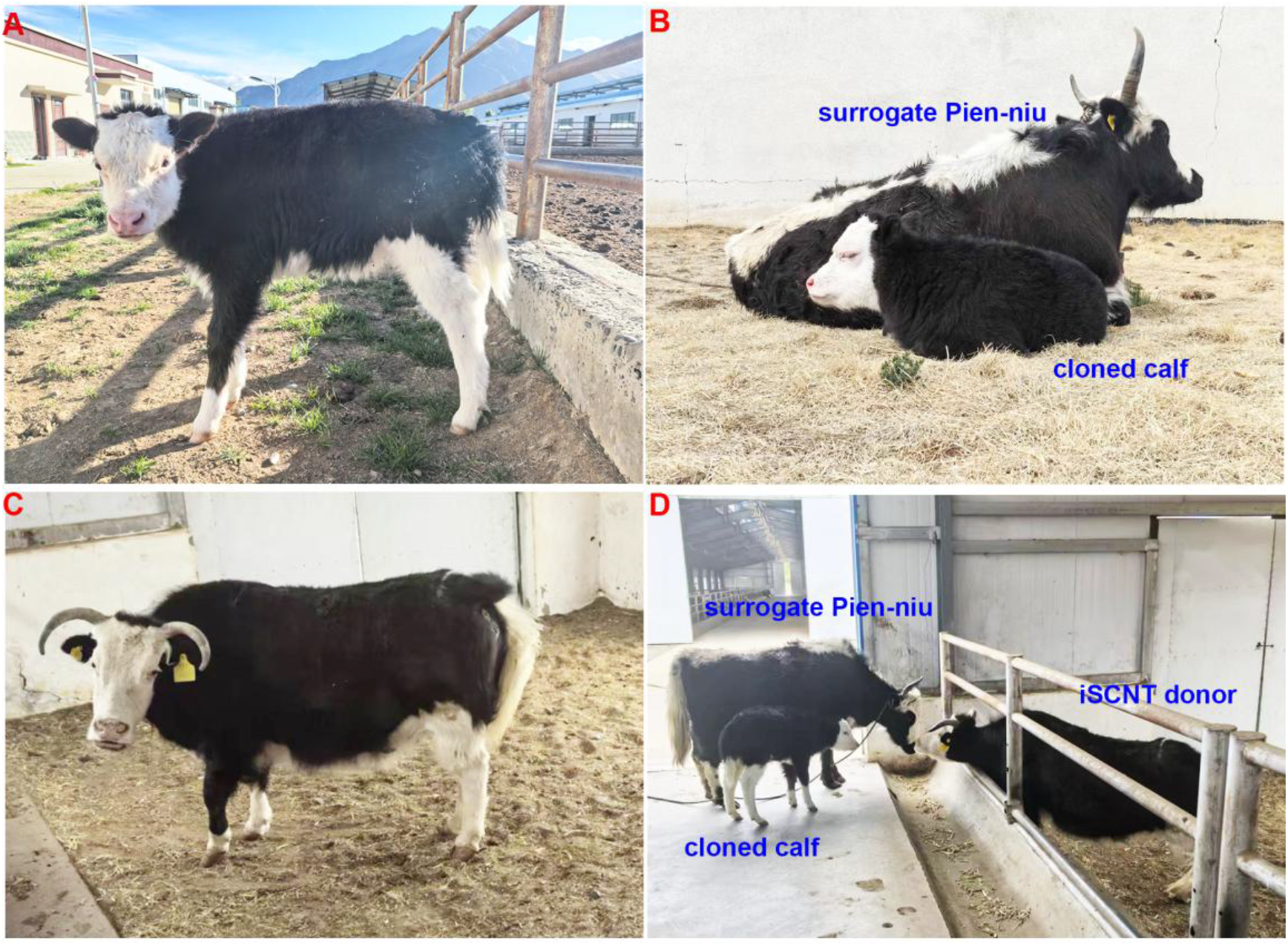
Production of cloned Pien-niu in Lasa Xizang. (A) The cloned calf derived from interspecies somatic cell nuclear transfer (iSCNT) embryo using Pien-niu ear fibroblast cell as donor and cattle oocyte as recipient. (B) The comparison of cloned calf and surrogate Pien-niu. (C) The Pien-niu provided ear fibroblast cell using as iSCNT donor. (D) The picture of cloned calf, iSCNT donor cell provider and surrogate Pien-niu.

### 2.2 Birth of cloned Pien-niu

The Pien-niu SCNT embryo transfer was performed at Qushui Experimental Station. In the present study, 9 female Pien-niu received spontaneous oestrus treatment were selected as SCNT embryo recipients. One well-developed SCNT blastocyst was transferred into each uterine horn. Pregnancy was confirmed via ultrasonography (Figure 2B), and 3 surrogated Pien-niu showed positive outcome, reaching a pregnancy rate of 30%. On May 12, 2025, a live cloned Pien-niu calf (Figure 1A) was delivered via cesarean section. Morphological analysis revealed that the cloned calf exhibited high phenotypic similarity (coat color, body conformation) to the nuclear donor Pien-niu, while showing distinct differences from the surrogate Pien-niu. Results of short tandem repeat analysis further revealed that the microsatellite loci of cloned calf were completely matched with that of donor fibroblast cells (Table 2, Figure 2). At its 60 day-old (July 10, 2025), the cloned Pien-niu calf is still in good health, and its weight has increased from 25.9 kg at birth to 71 kg (Figure 3), possibly indicating that the *in situ* conservation of Pien-niu at ultra-high-altitude region can be achieved via iSCNT.

**Table 2.** Analysis of short tandem repeat loci.

| Locus | Cloned calf | Donor cell | Surrogate cow |
| --- | --- | --- | --- |
| TGLA227 | 9/15 | 9/15 | 9/13 |
| SPS115 | 21/25 | 21/25 | 16/25 |
| ETH225 | 22/30 | 22/30 | 21/22 |
| INRA023 | 17/19 | 17/19 | 17/18 |
| ETH3 | 23/23 | 23/23 | 26/26 |
| BM2113 | 18/20 | 18/20 | 20/20 |
| TGLA122 | 18/22 | 18/22 | 18/22 |
| BM1824 | 13/17 | 13/17 | 17/18 |
| TGLA53 | 20/28 | 20/28 | 25/32 |
| SPS113 | 17/18 | 17/18 | 15/21 |
| CSRM60 | 19/21 | 19/21 | 17/22 |
| HAUT27 | 12/20 | 12/20 | 11/16 |
| ETH10 | 15/20 | 15/20 | 17/18 |
| ILSTS006 | 11/19 | 11/19 | 10/16 |
| MGTG4B | 6/17 | 6/17 | 6/21 |
| RM067 | 11/13 | 11/13 | 11/16 |
| BM1818 | 15/17 | 15/17 | 15/18 |
| TGLA126 | 13/21 | 13/21 | 16/21 |
| INRA063 | 11/13 | 11/13 | 8/10 |
| CSSM66 | 17/18 | 17/18 | 14/19 |

**Figure 2.**
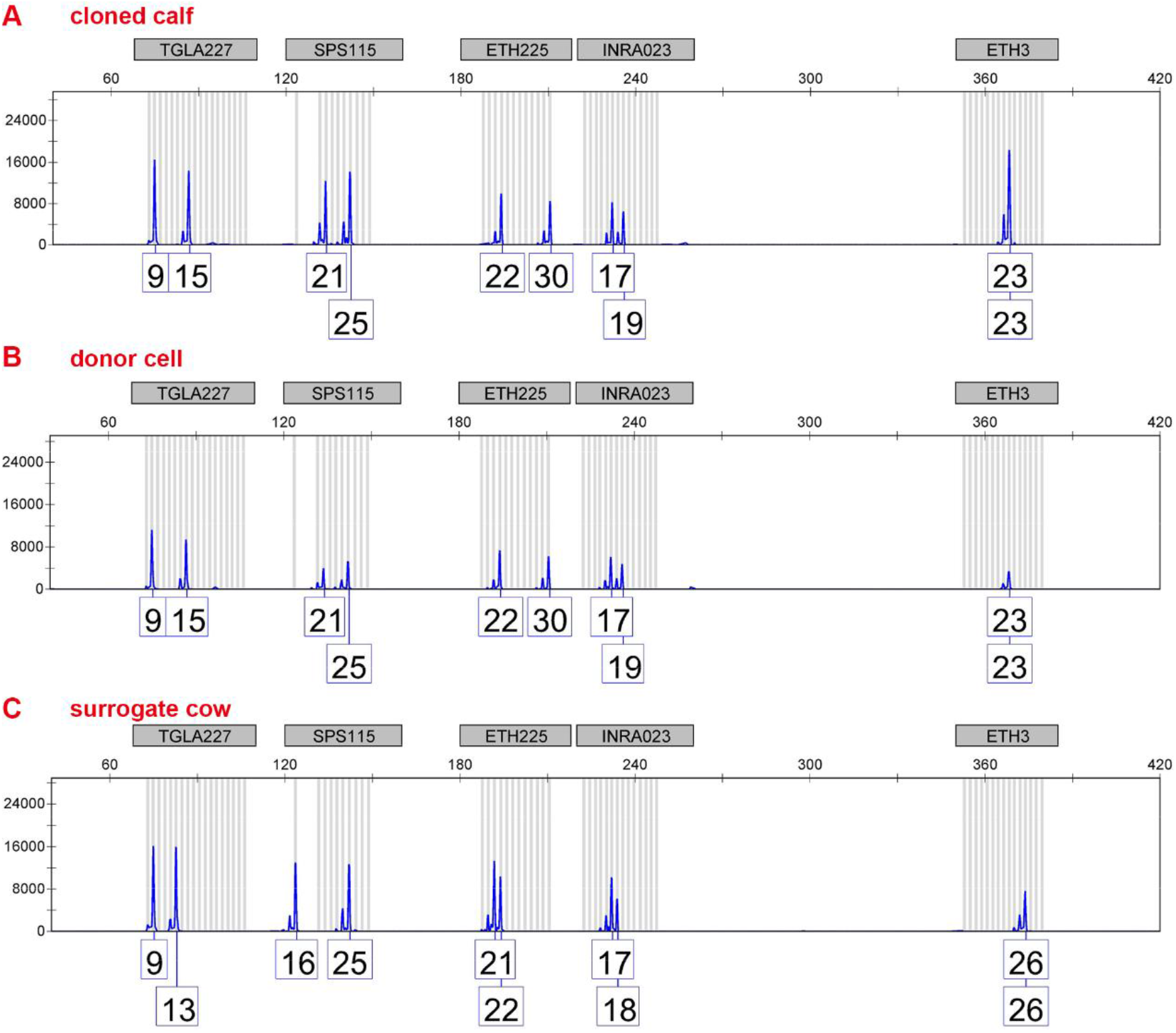
Electrophoresis peak map of the detection for some representative STR loci in iSCNT cloned calf (A), donor cell (B) and surrogate cow (C).

**Figure 3.**
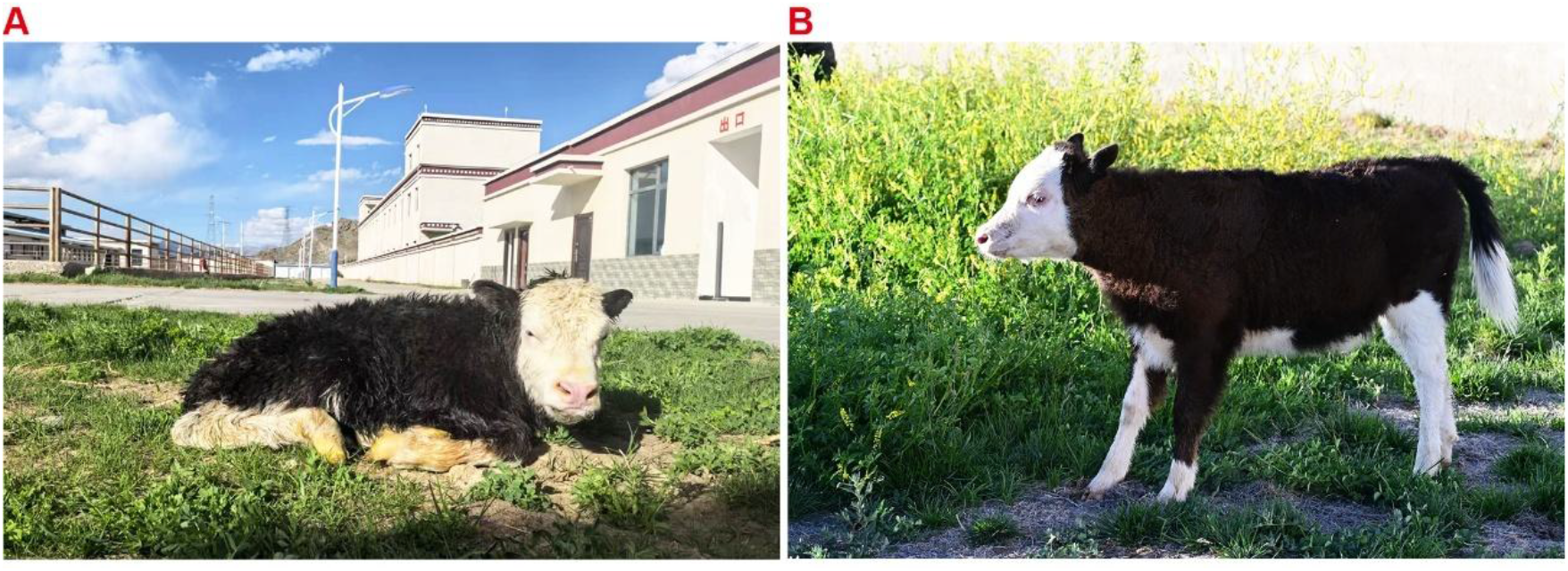
Appearance of the cloned Pine-niu at its first 60 days. (A) The picture was taken at its birth on May 12, 2025, and it weighed 25.9 Kg. (B) The picture was taken at its 60 day-old on July 10, 2025, and it weighed 71 Kg.

## 3. Discussion

The first production of cloned Pien-niu in Qinghai-Xizang Plateau announced the successfully overcoming the challenges of extreme environments to the SCNT technology. Previous studies revealed that when iSCNT embryos—constructed using ear skin fibroblasts from yaks (Bos grunniens) as donor cells—were transferred into 108 cattle recipients, only two pregnancies were achieved, neither of which progressed beyond 120 days of gestation ^11^. In addition, the provider Pien-niu of iSCNT donor cells and surrogate Pien-niu for iSCNT reconstructed embryos are all maintained at the same farm, Qushui Experimental Station, announcing the breakthrough in the *in situ* conservation at ultra-high-altitude region.

Realizing the goal of cloning Pien-niu in high-altitude area would be much more difficult and meaningful. Some local breeds and wild animals are endangered due to climate change, habitat degradation and overgrazing. SCNT technology, particularly interspecies SCNT, offers a viable solution to rapidly establish biobanks and expand populations of superior germplasm or endangered individuals, thereby effectively preserving genetic diversity ^12^. Cloning animals through SCNT has been an unpredictable process which often resulted in high rates of embryonic failure, stillbirths, and postnatal deaths ^13^. The cloning efficacy in the present study still requires improvement. Beyond the unique environmental factors in the Qinghai-Xizang Plateau, several critical factors—including epigenetic barrier in donor cells, and nucleocytoplasmic and mitonuclear incompatibility in the iSCNT embryos—may significantly impede reprogramming efficiency..

Current studies predominantly focus on purebred models. As far as we known, mules are one of the few hybrid animals that have been successfully cloned ^14^. The phenotypes of interspecies hybrids arise not only from interactions between nuclear genes but are also significantly influenced by cytonuclear interactions, which plays a crucial regulatory role in fetal and placental development^15^. Notably, hybrid organisms might present unique challenges for cloning technologies as conflicting parental epigenetic imprints may exacerbate reprogramming barriers, a critical scientific issue that warrants urgent investigation in future research.

## 4. Materials and methods

### 4.1 Animals

All experimental procedures were approved in advance by the Animal Welfare and Research Ethics Committee at the Institute of Animal Sciences, Chinese Academy of Agricultural Sciences (Approval Number: IAS2024-162).

### 4.2 Establishment of ear fibroblast cells

Ear tissues were collected from an adult female Pien-niu or cattle. The epidermal hair at ear edge was removed thoroughly with a sterile surgical knife. The tissue was washed with 75% alcohol, and then rinsed using phosphate buffered saline (PBS). Subsequently, the tissue was further cut into small pieces with surgical scissors, and digested in 0.5% trypsin solution in shaking incubator at 37°C for 35 min. After the digestion being terminated, the sample was centrifuged and resuspended in cell culture medium, and transferred into a 10 cm culture dish. The dish was then placed in a 5% CO_2_, 38.5°C incubator. When the cells grow into 80–90% confluence, they were digested using 0.5% trypsin and passaged.

### 4.3 *In vitro* maturation of oocytes

Ovaries were collected from a nearby slaughterhouse, washed with physiological saline solution, placed in a thermos flask, and then transported to the laboratory within 3 h. After being washed with physiological saline solution 3 times, 2-6 mm follicles were selected to aspirate the follicular contents using a 20 mL syringe. Cumulus oocyte complexes (COCs) with at least three uniform layers of cumulus cells were picked under the microscope. The COCs were cultured in mature medium containing 9.5 mg/mL M199, 1 mmol/L sodium pyruvate, 2.5 mmol/L L-glutamine, 10 ng/mL epidermal growth factor, 1 μg/mL 17β-estradiol, 10% fetal bovine serum, and 100 IU/mL penicillin for 16-18 h.

### 4.4 Somatic cell nuclear transfer (SCNT)

The maturated COCs were transferred into a microtube containing collagenase and pipetted repeatedly about 60 times. Denuded oocytes were placed in manipulation medium containing 5 mg/mL cytochalasin B (CB) and transferred to a micromanipulator. The oocytes were held by a holding needle, and a part of the ooplasm near the first polar body was aspirated by an injection needle. A donor ear fibroblast cell of 15 to 20 μm in diameter was aspirated by the injection needle and injected under the zona pellucida of the denuded oocyte. The donor cell-oocyte was electrically fused in a fusion medium, and transferred to M19 containing 10% FBS for culture after fusion for 2h. The successfully fused reconstructed embryos were treated in a medium 5 µmol/L ionomycin for 5 min, and then cultured in a medium containing 1.9 mmol/L 6-DMAP for 4 h. The activated reconstructed embryos were transferred to embryo culture medium and further cultured to develop into blastocysts.

### 4.5 Embryo transfer

Female Pien-niu with a healthy reproductive system, short calving intervals, and no history of dystocia were selected for synchronous estrus induction. Well-developed blastocysts were selected to be surgically transferred to the uterus of the surrogate Pien-niu at Qushui Experimental Station on the fifth day after estrus was observed. Pregnancy was detected by B-ultrasound on 30 days after embryo transfer.

### 4.6 Short tandem repeats (STR) analysis

Genomic DNA were extracted from the ear tissues of cloned calf and its corresponding surrogate cow, and the corresponding iSCNT donor cells for the cloned calf. Multiplex fluorescent PCR was performed using STR gene locus-specific primers. Capillary electrophoresis was used to separate the PCR amplification products, and the fluorescent signal was collected and analyzed. By comparing with the known standard for gene typing, the allele length of each STR locus was determined.

## Acknowledgement

This work was supported by the National Key R&D Program of China (2024YFD12007000), Key R&D Program of Xizang (CGZH2023000257), Innovation Program of Chinese Academy of Agricultural Sciences (CAAS-CSAB-202402), and the Open Project of State Key Laboratory of Animal Biotech Breeding (No. 2025SKLAB6-13).

## Competing interests

The authors declare no competing interests.

## Author contributions

D.Y., W.B. and Y.H. provided the conceptualization, funding acquisition, writing review and supervision. D.Y., Q.N., L.C., S.G., K.Y., Y.Z. and J.W. conducted investigation and formal analysis. D.Y. and Y.H. conducted the literature research and wrote the original draft. All authors have read and approved to publish the article.

## Notes

### Competing Interest Statement

The authors have declared no competing interest.

